# Roles of G1 cyclins in the temporal organization of yeast cell cycle - a transcriptome-wide analysis

**DOI:** 10.1101/287573

**Authors:** Lotte Teufel, Katja Tummler, Max Flöttmann, Andreas Herrmann, Naama Barkai, Edda Klipp

## Abstract

Oscillating gene expression is crucial for correct timing and progression through cell cycle. In *Saccharomyces cerevisiae*, G1 cyclins Cln1-3 are essential drivers of the cell cycle and have an important role for temporal fine-tuning. We measured time-resolved transcriptome-wide gene expression for wild type and cyclin single and double knockouts over cell cycle with and without osmotic stress. Clustering of expression profiles, peak-time detection of oscillating genes, integration with transcription factor network dynamics, and assignment to cell cycle phases allowed us to quantify the effect of genetic or stress perturbations on the duration of cell cycle phases. Cln1 and Cln2 showed functional differences, especially affecting later phases. Deletion of Cln3 led to a delay of START followed by normal progression through later phases. Our data and network analysis suggest mutual effects of cyclins with the transcriptional regulators SBF and MBF.

## Introduction

Eukaryotic cell cycle is a highly ordered process, which can be divided into four distinct phases during which a specific set of events take place: Cell growth (G1 and G2 phase), duplication and segregation of DNA (S phase) and the division of the nucleus (M phase), finally leading to cytokinesis. To enable and control the progression through the cell cycle, a subset of genes is transcribed in an oscillating pattern. Amongst those, cyclins are key regulatory proteins, which trigger all fundamental events of the cell cycle (Haase & Reed, 1999; Lu & Cross, 2010). Cyclins activate cyclin dependent kinases (CDKs), leading to phosphorylation of specific target proteins, which among other things, initialize the expression of the next wave of oscillating genes (for simplification we will refer to the cyclin-CDK complexes just by the name of their cyclins). The cyclins are functionally conserved across many species including mammals (Harashima *et al*, 2013), which makes understanding their functions even more relevant.

During G1 phase, the initial part of the cell cycle, three cyclins in *Saccharomyces cerevisiae* are necessary to successfully start the first regulatory events of the cell cycle: Cln1, Cln2 and Cln3. These G1 cyclins have been extensively studied and were found to fulfill specific functions but also to be able to partly compensate for each other in knockout studies. However, loss of all three cyclins at once is lethal for the yeast cells (Richardson *et al*, 1989), highlighting their importance for controlling cell cycle progression.

Cln3 is the first cyclin expressed during G1 phase, which, by inactivating the transcriptional repressor Whi5, is responsible for cell size control and the initial expression of the G1/S regulon. Two hypotheses have been proposed for the mechanism of the Cln3-Whi5 interaction: Both the retention of Cln3 in the endoplasmic reticulum with release in late G1 phase (Vergés *et al*, 2007; Wang *et al*, 2004) and the size-dependent dilution of Whi5 up to a critical threshold (Schmoller *et al*, 2015) can explain the Cln3-mediated release of Whi5 repression at the right time to initialize the transcription of genes required for the G1/S transition. In both cases, once Whi5 is sufficiently phosphorylated by Cln3, it is excluded from the nucleus and two transcriptional complexes, MBF (MluI Cell Cycle Box [MCB] Binding Factor), consisting of Swi6 and Mbp1, and SBF (Swi4/6 cell cycle box [SCB] Bining Factor), a heterodimer of Swi6 and Swi4 (Andrews & Herskowitz, 1989; Koch *et al*, 1993), can trigger the expression of genes in the G1/S regulon. However, the detailed wiring of this phase of the cell cycle network is still debated (for references see Figure 1).

**Figure 1:**
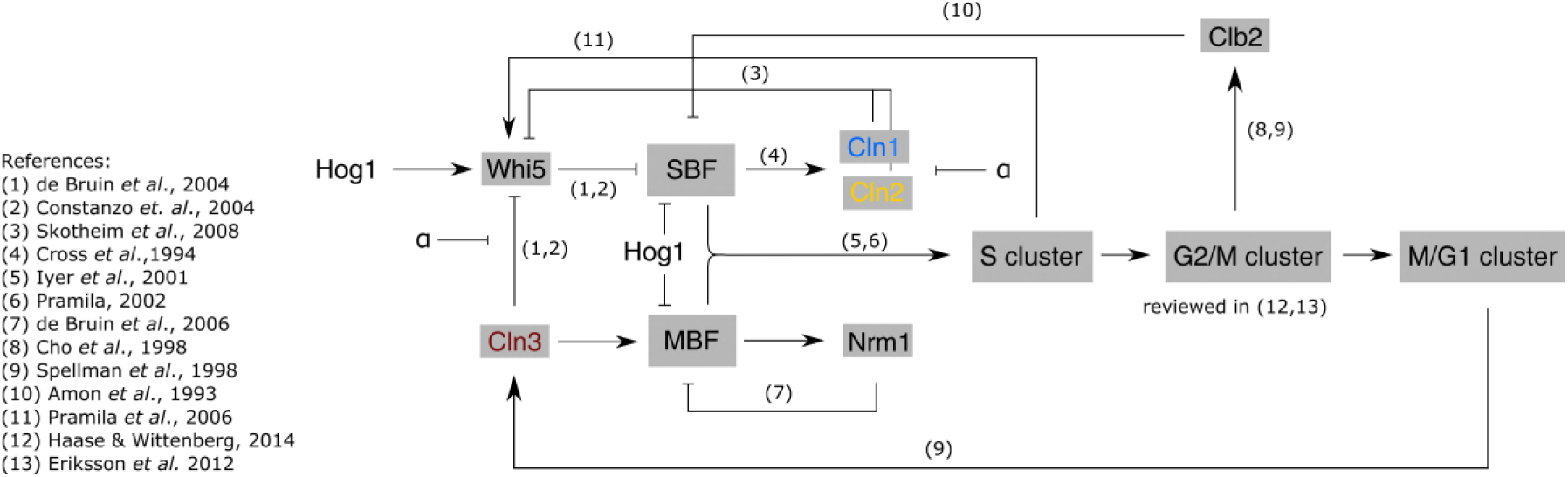
**Wiring diagram of the cell cycle**, based on references and results of this work. Main mechanisms of oscillating gene activation and inhibition are represented. G1 cyclins Cln1, Cln2 and Cln3 are shown in blue, yellow and red, respectively. G1 regulon (MBF/SBF) activation is shown in detail, activation of the following gene clusters (S, G2/M and M/G1 cluster) are only shown as an schematic overview. Activation is represented as arrows and inhibition as a bar-headed arrow. In addition, the effects of α-factor and osmotic stress are depicted. Related publications for each regulatory edge are shown as numbers next to the arrows.

The cyclins Cln1 and Cln2 are expressed in the G1/S regulon. They share the same wiring in the cell cycle network and are structurally highly similar (Queralt & Igual, 2004), which is why they are usually considered to carry out the same functions. In a positive feedback loop, both cyclins contribute to further phosphorylation of Whi5, thereby increasing the activation of MBF and SBF controlled genes and also their own expression (Skotheim *et al*, 2008). Besides this self-enhancement, Cln1 and Cln2 phosphorylate further targets, such as the S-phase inhibitor Sic1 (Schneider *et al*, 1996; Verma *et al*, 1997), leading to its degradation and a subsequent entry into S-phase in the continuing cell cycle.

This transition from G1 to S phase is called START and marks a point of no return. Hence, if a cell overcomes this checkpoint and commits to entering S-phase, it has to progress through the entire cell cycle. Accordingly, this checkpoint has to be tightly controlled to ensure that the cell is well prepared for a save passage to cell division. Many regulatory processes and pathways are therefore active before the checkpoint to control for both internal factors, such as cell size, genomic integrity or availability of storage compounds, and external conditions, such as available nutrients or environmental stresses. Many of these stresses induce a cell cycle arrest in G1 phase, which can be released to pass the checkpoint only after the stress has been counteracted by the cell. Well known examples are the response to an increase in external osmolarity or, in haploid cells of mating type *MAT***a**, the arrest due to the mating pheromone α-factor (Bucking-Throm *et al*, 1973). Both processes are also known to interact with the G1 cyclins: Osmotic stress, via the Hog1 signaling pathway, inhibits *CLN1* and *CLN2* expression (Escoté *et al*, 2004) and Cln3 activity (Bellí *et al*, 2001), whereas the pheromone pathway component Far1 is known to be a target of Cln1 and Cln2 (McKinney *et al*, 1993; Peter *et al*, 1993). The G1 cyclins are, therefore, key not only to normal cell cycle progression but also to the control of the cell cycle in stress scenarios.

In this work, we aim to dissect the specific functions of the three G1 cyclins in fine tuning of cell cycle timing. In particular, we are interested in understanding the contribution of each cyclin to organizing the passage through G1 phase and to identify effects on later cell cycle phases. A major question is thereby whether Cln1 and Cln2 are really fully redundant or whether we can identify specific influences on the cell cycle timing for each of them. To do so, we characterized the roles of the G1 cylins in organizing global oscillatory gene expression. We analyzed the transcriptome of wild type as well as single and double knockouts of Cln1, Cln2 and Cln3 to identify timing effects, such as overall delays or temporal shifts, in the expression patterns. For enhancing functional differences between Cln1 and Cln2, we additionally perturbed the cells with osmotic stress. Based on the results, we propose mechanistic links that could cause the observed timing effects depended on altered transcription factor activities.

We found that the deletion of the earliest cyclin Cln3 leads to a systematic shift of expression times, with an elongated G1 phase followed by a wild type like timing of the subsequent phases of the cell cycle. Loss of Cln2, on the other hand, results in an elongated G2 phase following a relatively conserved initial cell cycle. These timing effects correlate to changes in the activation of the G1 regulon triggered by MBF or SBF. Loosing Cln1 has a less dramatic effect than a Cln2 knockout, which only becomes evident when additional stresses are present: *cln1*Δ cells require longer times to exit from G1 phase after an osmotic shock and Cln1 alone (without Cln2 and Cln3) is not able to reliably release α-factor induced G1 arrest. Taken together our result show that the correct activation of the G1 regulon and, thereby, induction of oscillating gene expression, is strongly dependent on the G1 cyclins, each having specific roles in adjustment of cell cycle timing.

## Results

### Knockout of G1 cyclins alters cell size and transcriptional timing

To understand the mechanistic contribution of the major G1 cyclins Cln1-3 to the orchestration of the cell cycle, we analyzed the effect of their loss in single and double mutants compared to the wild type strain, BY4741, a haploid strain of mating type *MAT***a**. Those mutants are, as opposed to the triple mutant (Richardson *et al*, 1989), viable and allow for characterization of specific effects of the loss of each cyclin.

As was shown before (Soifer & Barkai, 2014; Talia *et al*, 2007), Cln genes act as regulators of cell size and morphology (Figure 2A). The loss of two out of three G1 cyclins causes the cells to increase in size compared to the wild type before division (Figure 2B). The loss of the initial cyclin Cln3 as single deletion already increases cell size comparable to double deletions.

**Figure 2:**
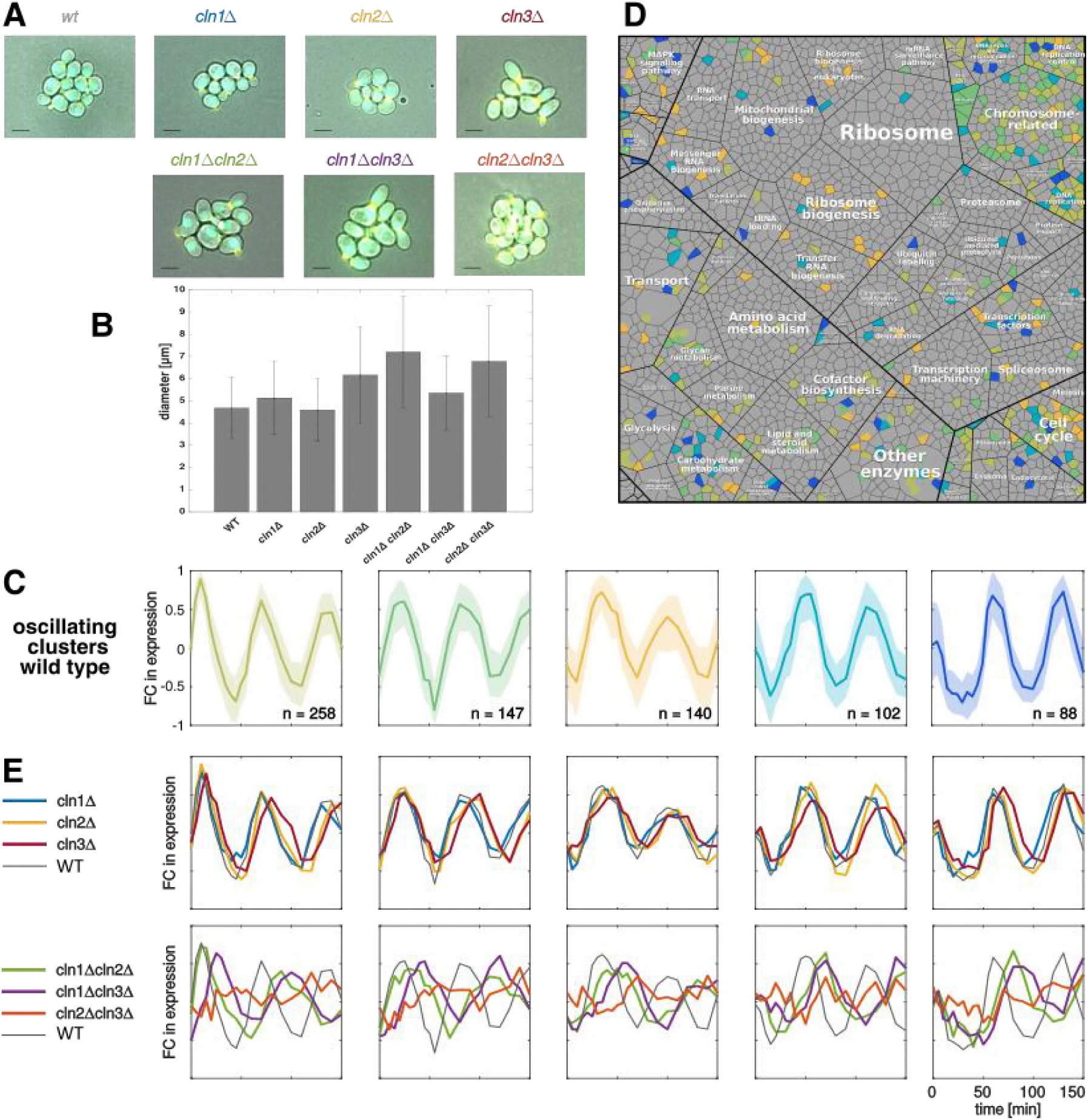
Characterization of cyclin deletion mutants. **(A)** Microscopic images of the used strains (overlay of brightfield and fluorescently labeled nucleus (Acs2-CFP) and bud neck (Cdc10-YFP)). Scale bar represents 4μm. **(B)** Cell sizes (diameter in μm) and standard deviation quantified by CASY® Cell Counter of wild type (WT) and knockouts. **(C)** Wild type expression of oscillating gene clusters, obtained by k-means clustering, sorted according to their peak times (fold change to mean of each gene, line represents mean of the cluster genes’ expression, shaded area 25% and 75% quantiles). **(D)** Functional classification of the oscillating genes based on a proteomap (Liebermeister *et al*, 2014). Each tile represents a gene, grouped according to its product’s function. Genes contained in the oscillating clusters in C are highlighted in the corresponding colors. **(E)** Mean expression of the oscillating gene clusters in the mutant strains in single mutants (upper panel) and double mutants (lower panel). For comparison, the wild type (WT) behavior is plotted in gray.

Changes in cell size can hint towards a deregulation of cell cycle timing, with altered cell cycle phase durations resulting in longer or shorter growth periods. We, therefore, set out to characterize the specific timing of cell cycle events. Progression through the cell cycle can be monitored by the periodic expression of genes in specific cell cycle phases (Eser *et al*, 2014; Santos *et al*, 2014; Cho *et al*, 1998; Spellman *et al*, 1998). Accordingly, we performed time-resolved RNA sequencing on the mutant strains following the release from synchronization with α-factor. To identify genes that showed an oscillatory behavior, we applied k-means clustering on the wild type gene expression trajectories (Figure 2C and Supplementary S.1). Oscillating genes grouped in 5 clusters with different peak times throughout the cell cycle. The wild type, thereby, showed two complete cell cycle periods within 150 minutes. A functional classification of the genes in the oscillating clusters showed that they take part in a wide variety of cellular processes (based on a generic proteomap (Liebermeister *et al*, 2014), Figure 2D). GO term analysis (Cherry *et al*, 2012; Ashburner *et al*, 2000) further revealed ordered timing of cell cycle functionalities for the wild type: Genes for DNA replication and chromosome organization (cluster 1) are followed by genes involved in chromosome segregation, chromatin assembly and microtubuli organization (cluster 2), subsequently in nuclear division and mitotic exit (cluster 4) and finally by genes required for the next cell cycle such as G1/S transition genes (cluster 5, for detailed results see Supplementary File 1).

The well characterized oscillatory gene subset in the wild type could then be used to examine the transcriptional changes occurring in the mutant strains (Figure 2E). The oscillating behavior of the gene clusters was conserved in the mutants except for the *cln2*Δ*cln3*Δ strain that did not show any oscillatory gene expression (lower panel, orange line). All other strains’ gene expression still oscillated, but showed timing effects of different magnitudes. While in the single knockouts we found only moderate changes in period and timing of the oscillations (upper panel), we observed strong delays and relative shifts in the double mutants (lower panel). Qualitatively, mutants lacking Cln3 showed a delayed expression in all 5 clusters, arguing for a delayed onset of the cell cycle, whereas the loss of Cln2, alone or in combination with Cln1, only delays the expression of later clusters (2-5), hinting towards a deregulation of individual cell cycle phase lengths.

### Cln mutants maintain transcriptional oscillations but not their timing

The clustering analysis revealed a qualitative overview of the implications of G1 cyclin loss. Quantifying these effects on the level of individual genes then allowed us to systematically characterize specific timing effects of each mutant, such as extensions of time intervals of each phase within the cell cycle.

Based on the expression data, we estimated the peak time φ of all measured genes with the help of the MoPS algorithm (Eser *et al*, 2014), (Figure 3A). The peak time estimates have a higher time resolution than our sampling intervals, such that we could also estimate peak times that lie between two probed time points. The algorithm furthermore provides a scoring of periodicity (see methods and Supplementary S.2) as well as an estimate for the period λ of the oscillations (Figure 3A), which corresponds to the cell cycle length. The median cell cycle duration of the so identified subset of oscillating genes (Supplementary File 2, Supplementary S.3) was 63.5 minutes in the wild type (Figure 3B). The single deletion mutants only showed a slightly changed cell cycle duration with a more prominent elongation effect of 4.5 min and 9.5 min in *cln2*Δ and *cln3*Δ, respectively. The double deletions *cln1*Δ*cln2*Δ and *cln1*Δ*cln3*Δ showed a clear extension by 22 min and 20 min of the cell cycle compared to the wild type (Figure 3B) in accordance with their increased cell size. Due to the loss of oscillating gene expression, cell cycle duration could not be estimated with this method in the *cln2*Δ*cln3*Δ strain.

**Figure 3:**
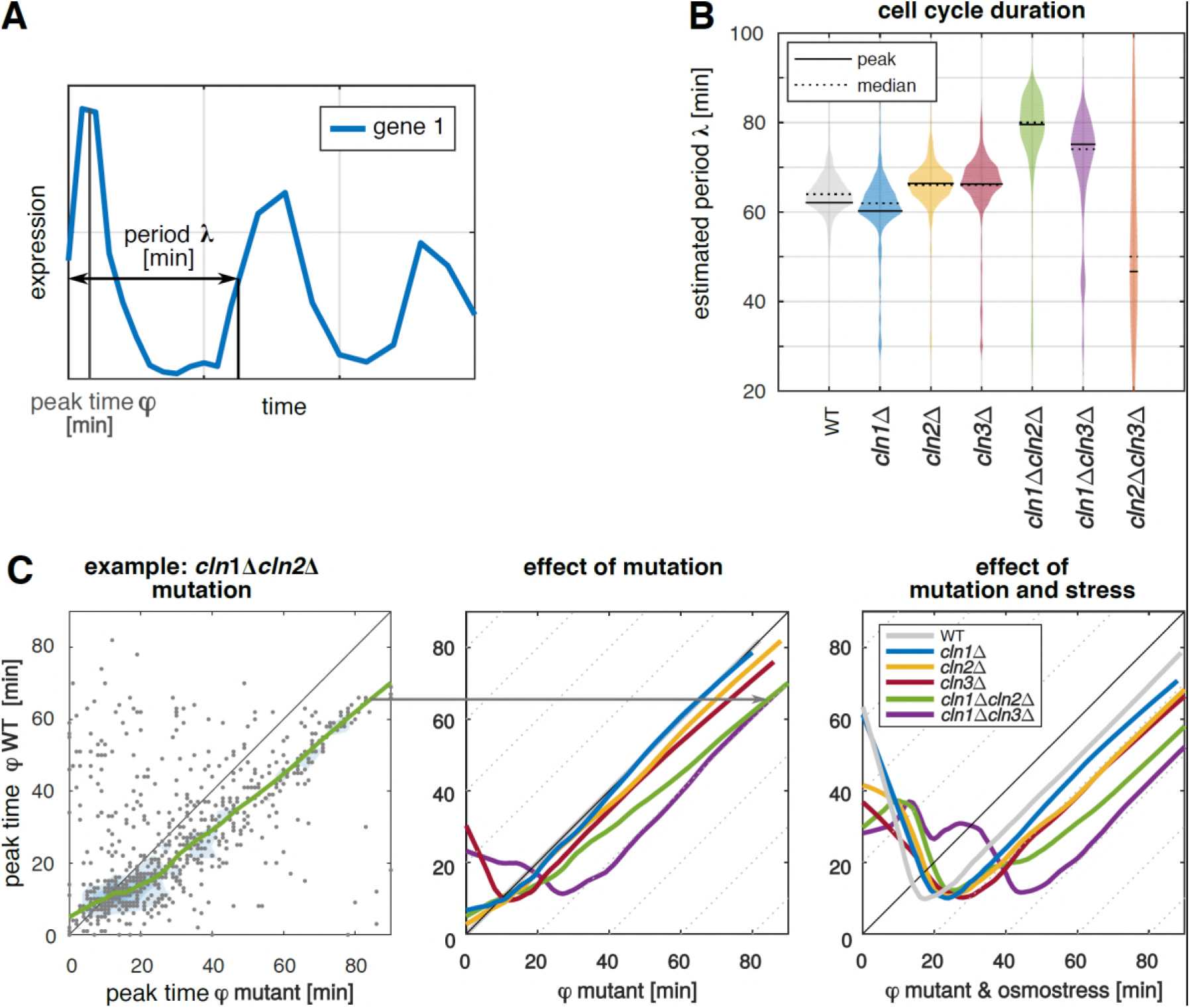
Quantitative peak time analysis. **(A)** Schematic representation of the oscillation properties of a gene, as estimated by MoPS algorithm (Eser *et al*, 2014). **(B)** Estimated period λ, corresponding to the cell cycle duration, of wild type (WT) and mutants (solid line represents most frequent period and dotted line represents median of period) as estimated using the MoPS algorithm (Eser *et al*, 2014). **(C)** Shift in estimated peak times φ. Left: as example cln1Δcln2Δ is shown. All genes contained in the oscillating clusters for mutants w.r.t. the wild type expression (y-axis) are presented. Each gray dot represents one gene, colored lines are lowess smoothed curves. While genes occurring at the bisecting line have the same peak time in mutant and wild type, genes below the bisecting line have higher peak times in the mutant. Middle: Effect of mutations on peak time φ w.r.t. wild type, lowess smoothed curves for all mutants except of cln2Δcln3Δ are shown. Right: Combined effect of mutation and osmotic stress compared to unstressed wild type.

The estimated peak times of the individual oscillating genes were then used to systematically analyze quantitative differences in the expression timing. For each mutant, peak times were compared to the wild type estimates in a two-dimensional scatterplot (as an example *cln1*Δ*cln2*Δ is shown in Figure 3C, left). We applied lowess smoothing to better visualize the general behavior of the entire group of oscillating genes. As genes occurring close to the bisecting line have a conserved peak time in mutant and wild type, we can use the slope of the smoothed curve to identify delays in specific time intervals (characterized by slopes smaller than one). Shifted curves with a conserved slope around one then indicate a overall delay with a conserved peak timing, for example caused by a delayed onset of the cell cycle.

With all peak times gathering around the bisecting line, no effect of the loss of Cln1 alone on the overall cell cycle timing was evident (Figure 3C, middle, blue line). In contrast, cells lacking Cln2 (as single deletion (yellow) or in combination with Cln1 deletion (green)) showed a conserved timing of the early peaks, but a decreased slope for later times – arguing for a slower progression through later cell cycle phases such as S and G2 phase. In the *cln1*Δ*cln3*Δ double mutant (purple) and, slightly less prominent, in the *cln3*Δ single deletion (red), we observed that the overall sequence of the peak times was retained (slope close to one) with a delay of around 5 and 20 minutes, respectively. However, a subset of genes shifts their peak time backwards to the first third of the cell cycle, visible as a negative slope of the lowess curve. A thus elongated G1 phase, in which many genes show their peak expression, is a behavior often observed in cells that react to a stress stimulus.

Following this thought, we repeated the peak time analysis with cells that had been exposed to osmotic stress following their release from synchronization. Osmotic stress is among others known to arrest cells in the G1 phase of the cell cycle until the stress is counteracted (Maeda *et al*, 1994; Alexander *et al*, 2001; Bellí *et al*, 2001; Yaakov *et al*, 2003; Escoté *et al*, 2004). Arrest in later phases is neglected here, since cells were exposed to osmostress in early G1. Since also our target cyclins act primarily in the G1 phase, we hypothesized that an additional perturbation in this phase could reveal further aspects of their action. In the stress experiments, we observed a delay of oscillating gene expression for wild type and all mutants (Figure 3C, right). Also in the *cln1*Δ strain, which did not show changes in expression timing in the unstressed experiment, we could now observe a stronger delay than in the wild type (11.5 mins delay caused by osmotic stress in the wild type, 16.9 mins in *cln1*Δ), highlighting the role of Cln1 in reentering the cell cycle after osmostress.

In summary, the peak time analysis revealed specific characteristics of the mutant gene expression: The loss of Cln3 causes an overall delay in oscillating gene expression, while the loss of Cln2 results in a relative shift of peak events towards later cell cycle times. Furthermore, by means of stress experiments, we could demonstrate that Cln1 and Cln2 are functionally non-redundant, whereby loss of Cln1 causes deregulation of transcriptional timing under osmostress.

### The network of periodically acting transcription factors reveals phase specific timing effects

The observed timing differences already contributed to the understanding of Cln functionality, but so far lacked mechanistic explanation. Our RNAseq data provided a characterization of the timing of gene expression, which is usually the result of the action of transcription factors. We, therefore, analyzed the systemic differences in transcription factor action in the mutants to wild type in order to identify candidate factors responsible for the timing effects.

We utilized a generic transcription factor network (YEASTRACT (Teixeira *et al*, 2014), Supplementary file 3) and reduced it to the oscillating subnetwork of our interest. To do so, we used the MoPS algorithm (Eser *et al*, 2014) to calculated the periodicity scores of the mean trajectory of all differentially expressed targets of each transcription factor in the wild type dataset. We discarded all transcription factors with a target periodicity score smaller than 0.75 or with less than 3 target genes. In the resulting, fully oscillatory network (Figure 4A, Supplementary Table 1) many transcription factors are present that are known to be involved in the cell cycle. Consistent with the expected information flow in the network, most of the regulatory edges connecting the oscillating transcription factors are directed from early to later peaking genes (top to bottom in Figure 4), only some are in the reverse direction (*e.g.* from Ace1, Swi4, Ste12, Tec1). These backward regulations could either be explained by inhibitory function of the transcription factor or by regulation of early genes in the following cell cycle.

**Figure 4:**
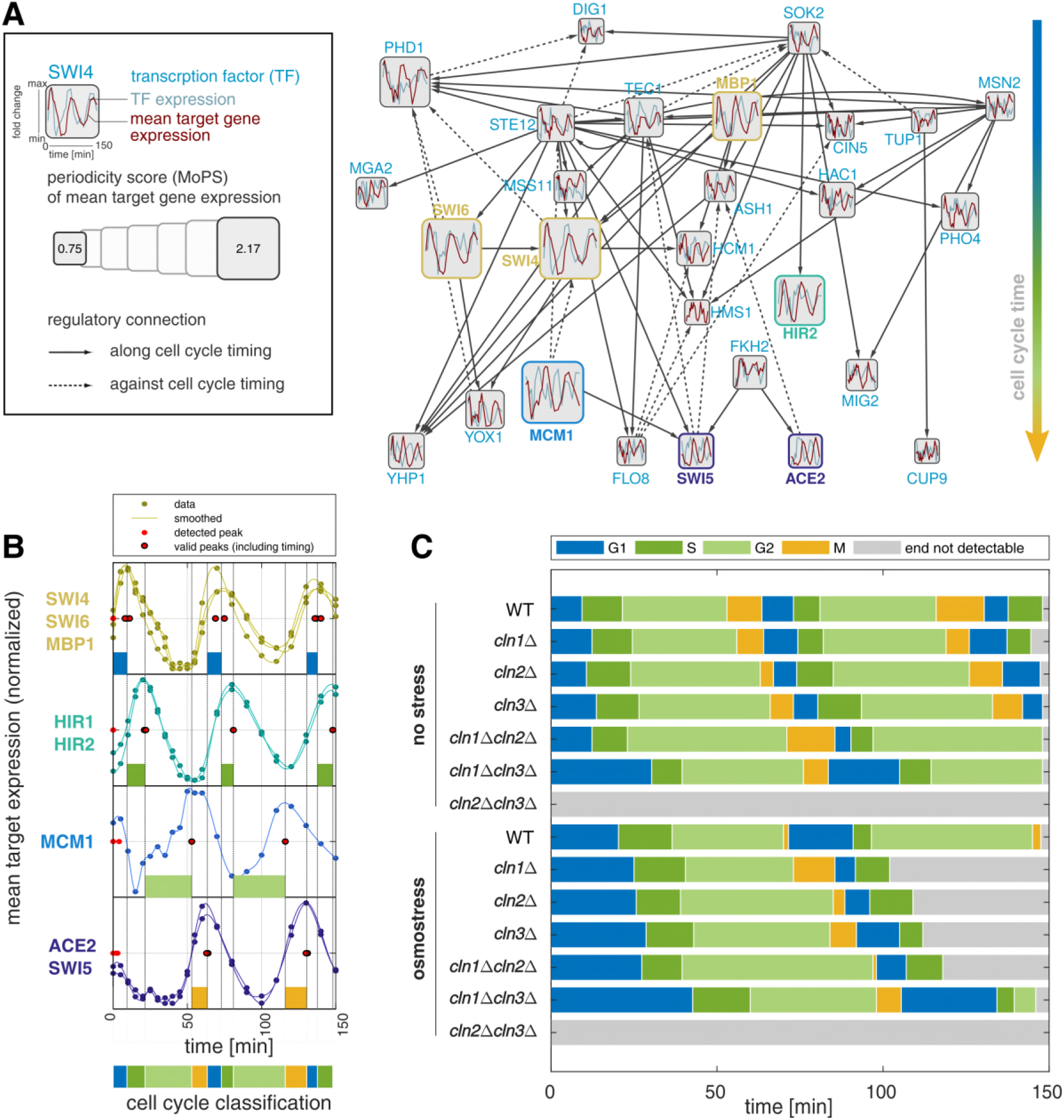
Cell cycle phase classification by active transcription factor (TF) network. **(A)** Summary of the oscillating transcription factor network in the wild type (WT). Vertical arrangement follows the target gene peak times (TFs with early peaking targets at the top). Regulatory edges that adhere to the cell cycle timing are shown as solid lines (start-point TF’s targets peak earlier than end-point targets) and as dashed lines otherwise. **(B)** Wild type target expression of “classification set” transcription factors used to define cell cycle phase durations, trajectories depict mean expression level of the targets (log fold change to mean). Phase transitions were defined according to the rules in Supplementary Table S.2. **(C)** The accordingly classified phases of all strains and conditions. See also Supplementary Figure S.4.

Based on this wild type regulatory network, we could identify transcription factors that showed a changed temporal expression in the mutants. To reduce complexity, we focused on the transcription factors with the highest periodicity score. Thereby, we selected factors representative for specific cell cycle times, which often corresponded to transcription factors known to be active at the transition between cell cycle phases. Our final “classification set” of transcription factors (Supplementary Table 2) consists of four groups, whose peak expression occurs at the transitions between G1/S, S/G2, G2/M and M/G1 phase (wild type example in Figure 4B, mutants in Supplementary Figure 4). From the classification set, we then estimated the cell cycle phase durations in all stressed and unstressed mutants (Figure 4C). It is important to notice that we detected the peak expression times here with a simple peak detection based on the smoothed target trajectories as opposed to using the MoPS peak time estimates. The reason for this is that especially for the stress experiments, but also due to the influence of the α-factor synchronization, the first measured cell cycle in our experiment can have different phase lengths than the second one. The MoPS algorithm (Eser *et al*, 2014), however, uses one fixed period for the entire time course. While this was not a problem in our previous analyses, which aimed at identifying oscillatory genes, we are now interested in the specific shifts in the phase timing.

Due to the lack of oscillations, no cell cycle phases could be assigned for *cln2*Δ*cln3*Δ, while we could identify all phases as well as specific timing effects for all other strains. All single deletions showed a slight elongation of G1 phase with the strongest effect in *cln3*Δ. The double deletions had stronger effects on the cell cycle timing. *cln1*Δ*cln3*Δ showed the longest G1 phase, which is without stress longer than the osmo-stressed wild type G1 phase. In contrast, not only G1 but also G2 phase was extended in the *cln1*Δ*cln2*Δ strain. It is known that cells grow especially in G1 phase and that a G1 arrest leads to bigger cells (Johnston *et al*, 1977; Skotheim *et al*, 2008; Turner *et al*, 2012). This is supported by our results showing that cells with extended G1 phase, like *cln3*Δ and the double deletion *cln1*Δ*cln3*Δ were significantly bigger than the wild type (Figure 2B). Additionally, our data showed that an increase in size is not only dependent on G1 duration but also on G2 duration. Demonstrated, for example, by size differences of *cln1*Δ and *cln1*Δ*cln2*Δ both having the same G1 phase (12.3 min) duration but the double deletion has a significant longer G2 phase (*cln1*Δ: 31.5 min, *cln1*Δ*cln2*Δ: 48 min), corresponding to bigger size of *cln1*Δ*cln2*Δ.

The addition of a high concentration of osmolytes arrests yeast cells in G1 phase, until cells finish their stress response and are once again ready to commit to a new cell cycle (Bellí *et al*, 2001; Escoté *et al*, 2004). This additional elongation of the G1 phase was still functional in the mutant strains: *cln2*Δ and *cln3*Δ were exiting the arrest later than the wild type and had an accordingly longer cell cycle. Also for the *cln1*Δ strain, we could see an extended cell cycle duration, mainly caused by elongated G1 and M phases. However, the longer M phase has to be handled with care, as the definition of the G2/M transition was often hampered by a plateauing expression of MCM1 targets. In the stress experiment, G2 phase is already extended when Cln2 alone is lost and, consistent with the unstressed case, requires an even longer time in the *cln1*Δ*cln2*Δ strain.

The individual phase lengths defined by the oscillating transcription factor network then allowed us to dissect the observed timing effects: Cells lacking Cln3 spend longer time in G1 phase, in which the expression triggered by the relevant transcription factors (SWI4, SWI6, MBP1) rises less rapidly than in the remaining mutants. In cells lacking Cln2, the shifted peak times occurred in an extended G2 phase, which was marked by an delayed expression of the targets of the later transcription factors MCM1 and ACE2/SWI5. The loss of Cln1 resulted in a less prominent but similar behavior.

### Loss of CLN3 synchronizes SBF and MBF target expression

In the Cln3 deletion mutants, we observed altered expression in the first group of transcription factors from our classification set. Those proteins are known to form the functional complexes MBF and SBF, which critically regulate the expression of the G1/S regulon and, thereby, facilitate the transition between G1 and S phases (Nasmyth & Dirick, 1991; Ferrezuelo *et al*, 2009; Eser *et al*, 2011; Orlando *et al*, 2008; Teixeira *et al*, 2014; Lee *et al*, 2002; Guo *et al*, 2013). Functionally, MBF targets are mostly involved in organizing DNA replication and adjunct processes while SBF targets control cell morphology (Lowndes Noel F., 1991; Igual *et al*, 1996; Iyer *et al*, 2001). Cln1-3 are involved in the regulation of MBF and SBF and, in the case of Cln1 and Cln2 also themselves part of the G1/S regulon (Wittenberg *et al*, 1990; Cross *et al*, 1994). MBF and SBF are regulated by slightly different mechanisms that share part of their actors. The explicit roles of the cyclins in this regulation, especially for MBF, are, however, still unknown.

Based on the high temporal resolution of our data, we could analyze the expression pattern of the two transcriptional complexes’ target genes in detail. We calculated two quantitative measures based on auto-correlations: (i) the delay between the expression of SBF and MBF target genes in each mutant (Figure 5 A) and (ii) the delay of MBF target gene expression in each mutant compared to the wild type expression (Figure 5B).

**Figure 5:**
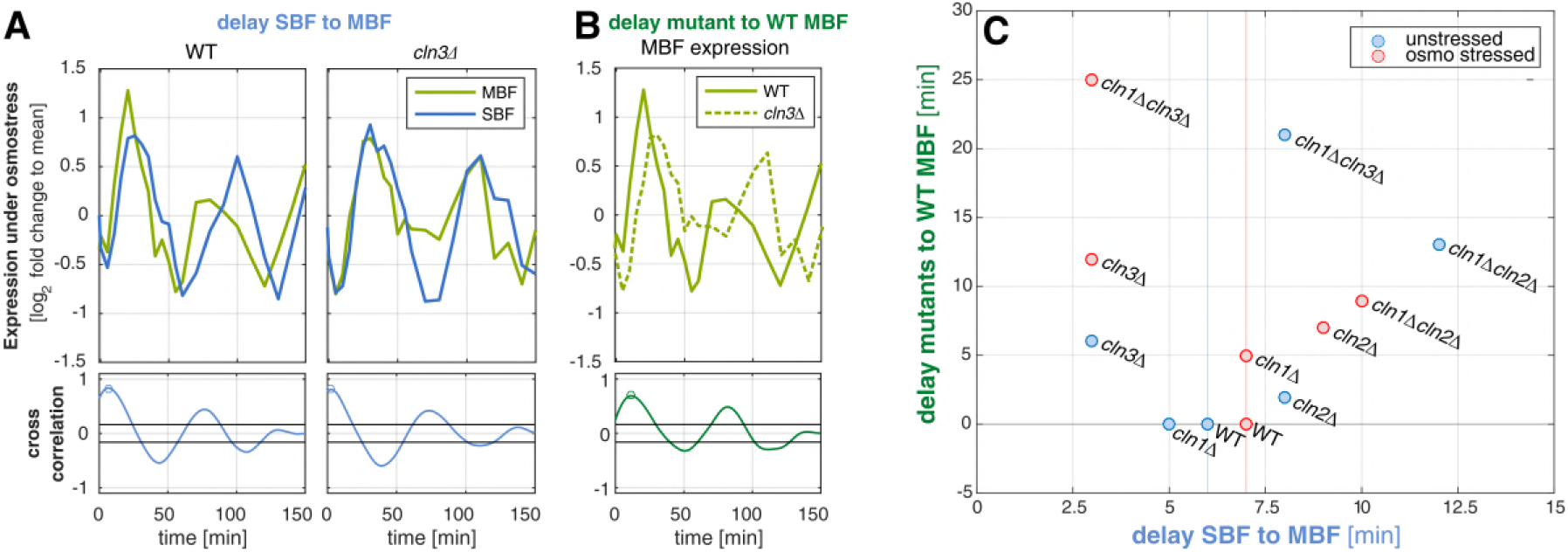
Detailed view on MBF and SBF target gene expression. **(A)** Delay between MBF (green line) and SBF (blue line) target expression, shown exemplary for wild type (WT) and cln3Δ. Upper part shows the mean expression of all target genes (normalized as log2 fold change to the temporal mean of each gene), lower part indicates the cross correlation of the two trajectories, the maximum is marked with a circle. **(B)** Delay of MBF target gene expression in a mutant (as example cln3Δ is shown, dotted line) to the wild type expression (solid line). Upper and lower panels as in A. **(C)** Summary of the measures exemplified in A and B for all mutants under both unstressed (blue dots) and osmostressed (red dots) conditions. The delay between SBF and MBF expression is shown on the x-axis, the delay of the mutant to the wild type MBF expression on the y-axis.

In α-factor synchronized cells, MBF targets are expressed some minutes before SBF targets (Eser *et al*, 2011). We observe the same behavior in the wild type, in which the SBF peak occurs around 6 minutes after the MBF peak. Also our mutant strains showed the same temporal order, but the delays between the two target cluster expressions were affected by the mutations (Figure 5C). SBF expression was delayed by 8 min and 12 min, relative to the expression of MBF target genes, in strains lacking Cln2 as single and double deletion together with Cln1, respectively. Contrarily, in cells lacking Cln3, MBF expression was delayed strongly compared to the wild type, such that it almost coincided with the expression of SBF target genes and the difference between the two clusters was shortened to around 3 minutes. Applying osmotic stress the observed behavior becomes even more pronounced. Because the definition of MBF and SBF target genes is not necessarily unique and also based on experimental evidence, we repeated the analysis based on further previously published target lists (Supplementary S.5, target lists in Supplementary File 4, (Teixeira *et al*, 2014; Eser *et al*, 2011; Macisaac *et al*, 2006; Ferrezuelo *et al*, 2009). The obtained results were qualitatively reproducible for all four different target lists, even though they showed differences in the absolute values for the calculated delays. Especially, the *cln1*Δ*cln3*Δ strain showed a stronger synchronization of SBF and MBF target expression in the sets obtained from the further sources. Focusing on the estimated cell cycle phase durations, in which the Cln3 mutant strains showed extended G1 phases, we could now link this to the delayed expression of the MBF gene cluster. Accordingly, Cln3 has to have a - direct or indirect - role in the activation of MBF, which has been proposed before. The observed elongation of the later G2 phase, on the other hand, occurred in the strains that show a relative delay of the SBF cluster to the MBF cluster.

### *cln2*Δ*cln3*Δ cells show defective release after α-factor synchronisation

In combination, our results outline a predominant influence of Cln3 and Cln2 on the timing of the early cell cycle phases. Consistently, but also unfortunately for further analysis, the *cln2*Δ*cln3*Δ strain did not show oscillations in gene expression (Figure 6A), indicative of a disruption of cell cycle timing. Nevertheless, the mutant is able to grow and divide in culture (Figure 2A) and to orchestrate an appropriate transcriptional response to osmotic stress (Supplementary Figure S.6).

**Figure 6:**
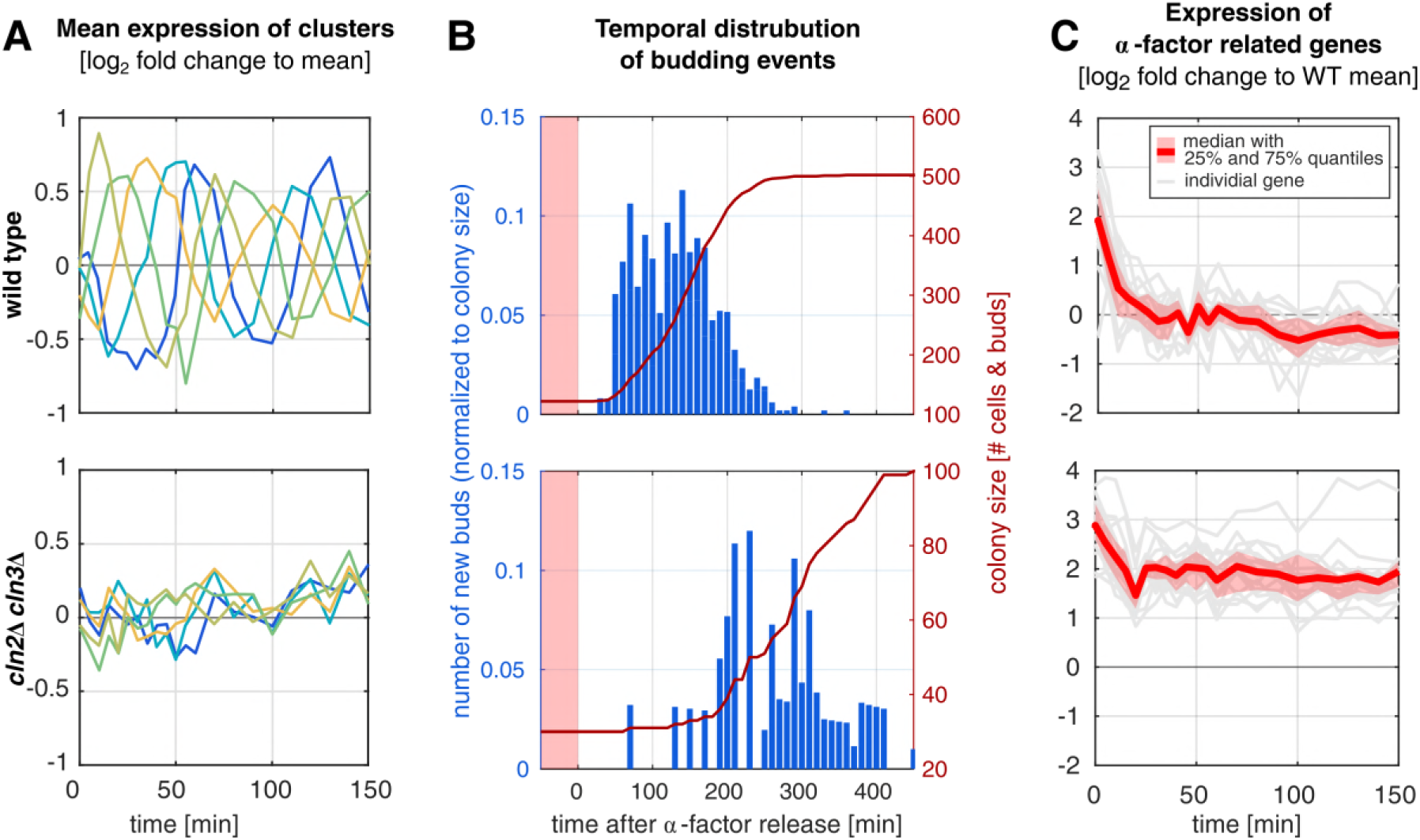
Absence of oscillations in the cln2Δcln3Δ strain is due to defective release from α-factor arrest. Comparison of wild type (top panel) and cln2Δcln3Δ (bottom panel). **(A)** Expression of genes in the oscillating clusters (same as in Figure 2 C and E). **(B)** Timing of bud appearance measured in a microfluidic setup with wash-out of α-factor at time point 0, after 180 minutes of synchronization in which no budding occurred in both strains. Several colonies were imaged every 10 minutes and newly occurring buds were counted for each image (blue histogram), red line represents colony size as total number of cells and buds. Budding was detected for two cell cycles after the release. **(C)** Group of α-factor induced genes showing higher expression in cln2Δcln3Δ than in wild type, red line represents mean expression with 25% and 75% quantiles, gray lines represent individual genes (see also Supplementary Table S.3).

We, therefore, hypothesized that possibly already the release from α-factor synchronization could be affected in the *cln2*Δ*cln3*Δ strain, resulting in an asynchronous re-entry to the cell cycle. In the RNAseq data we would then fail to detect oscillations because we are actually measuring an unsynchronized cell population.

To test this hypothesis, we followed α-factor release of the cells in a microfluidic device, recording the time points of budding events (Figure 6B). Indeed, first buds following a change to medium without α-factor occurred much later and during a longer time range in the *cln2*Δ*cln3*Δ strain than in the wild type (and also all other mutants, see Supplementary Figure S.7).

To understand the effect also on the transcriptional level, we could harness the results of the cluster analysis again. Besides the previously described oscillating clusters, we also found a set of genes whose expression is highest at the beginning of the experiment and decreases steadily afterwards. The cluster behavior was robust in all mutants, but showed the weakest decay in the *cln2*Δ*cln3*Δ strain (Supplementary Figure S.1, cluster 4). Based on functional enrichment of genes for conjugation, sexual reproduction, cell aggregation and cellular response to pheromones in this cluster (Supplementary File 1), we could characterize the behavior as a decaying response to α-factor arrest. Zooming in even further, we identified a group of 16 genes related to α-factor signaling and its cellular response (Supplementary Table 3), which remained at levels 2.5 - 7.7-fold higher in the *cln2*Δ*cln3*Δ strain than in the wild type (Figure 6C). We, therefore, concluded that the double mutant lacking both Cln2 and Cln3 fails to silence α-factor signaling in a coordinated fashion once the pheromone is removed from the medium. Individual cells, therefore, needed different times to exit the cell cycle arrest, which resulted in a smeared mRNA expression in the population that did not reveal oscillations. Other coordinated processes, which are largely independent of the cell cycle, can hence still be seen in the transcriptome such as here the response to osmotic stress. Another crucial point is that the defective release occurred in the *cln2*Δ*cln3*Δ but not the *cln1*Δ*cln3*Δ mutant, additionally highlighting the difference between Cln1 and Cln2 and their control over the timing of the cell cycle. As before, loss of Cln2 had much more dramatic consequences for the cells than loss of Cln1.

## Discussion

The G1 cyclins Cln1-3 are essential players in the initialization of the cell cycle (see schematic representation in Figure 1). Based on transcriptome-wide, time-resolved gene expression data in different Cln knockout strains, we showed functional differences between them and explored their contribution to the fine tuning of the cell cycle. Specifically, we showed that Cln1 and Cln2 have similar but clearly non-redundant functionalities, with Cln2 exerting stronger control over the cell cycle timing and influencing it beyond the initial G1 phase. Cln3, on the other hand, is known to have a distinct mechanism of action upstream of Cln1 and Cln2 activation. Consistently, we here characterized a strong and unique delay pattern in the *cln3*Δ mutant and proposed a MBF related mechanism for the observation. Based on the identified major roles of Cln2 and Cln3 in the start-up of the cell cycle, we could also explain the lack of transcriptional oscillations in a mutant lacking both genes due to the inability to exit from α-factor arrest in an ordered fashion.

### Cln1 and Cln2 are functionally semi-redundant

The two cyclins Cln1 and Cln2 are usually thought to have redundant function (Hadwiger *et al*, 1989; Benton *et al*, 1993; Haase *et al*, 2001; Peter *et al*, 1993), and in functional models of the cell cycle they are often lumped together in one species (*e.g.* (Chen *et al*, 2004; Li *et al*, 2004; Barberis *et al*, 2007; Barik *et al*, 2016). Experimentally, Cln1 and Cln2 show slight differences in expression timing (Eser *et al*, 2011), degradation pattern (Quilis & Igual, 2017) and nuclear accumulation (Quilis & Igual, 2012; Queralt & Igual, 2004), questioning a full redundancy of the two cyclins.

We here showed that loss of Cln2 had a more dramatic effect on the cell cycle timing than loss of Cln1. Cells lacking Cln2 showed a stronger shift in gene expression peak times (Figure 3), a more prominent delay of SBF to MBF target gene expression (Figure 5) compared to the *cln1*Δ mutant as well as a defective release from α-factor arrest in combination with the knock-out of Cln3.

Both cyclins are part of a self-regulating positive feedback loop (Figure 1). The found functional differences finally allowed for the hypotheses that (i) repression of the transcriptional repressor Whi5 by Cln2 is stronger than by Cln1 and that (ii) the pheromone response is predominantly interacting with Cln2 (see below).

### Loss of the positive G1 feedback increases G2 phase duration, concurrent with delayed SBF action

A surprising finding from our data was the elongation of the G2 phase in cells lacking Cln2 (Figure 4). In these mutants, we also saw a relative delay of SBF to MBF target gene expression. When the positive feedback loop via Cln2 is lost or reduced in its activity, the repression of SBF by Whi5 can only be relieved by the action of Cln3 (compare Figure 1), which seems to be far slower, causing a delayed expression of SBF targets genes. A potential regulation of Swi4, which is part of the SBF complex, by MBF (Harris *et al*, 2013) seems to only partly compensate for the action of the feedback.

The delay in SBF expression could affect the length of later phases of the cell cycle, as observed here for the G2 phase, via indirect links. For example, the transcription factor NDD1 is under the control of SBF (Macisaac *et al*, 2006). NDD1 activates the transcription of a gene cluster including CLB2 at the G2/M transition, which exerts a repression feedback on SBF (Amon *et al*, 1993; Koch *et al*, 1993; Siegmund & Nasmyth, 1996).

### Loss of Cln3 delays start of cell cycle and MBF activation

Amongst the single mutants, *cln3*Δ showed the strongest phenotype, including increased cell size, cell cycle duration, deregulation of oscillating gene peak times and number of differentially expressed genes (Supplementary S.8). The slope-conserving shift of all peak times in that mutant (Figure 3) indicated an overall delay of the onset of the cell cycle, consistent with an elongated G1 phase (Figure 4). Furthermore, we here characterized a delay of MBF target gene expression in mutants lacking Cln3 (Figure 5) that correlated with the delayed onset of the cell cycle. That hints to a functional connection between Cln3 and MBF activation.

From previous literature, the detailed mechanism of MBF activation is not yet known. It has been proposed that MBF de-repression at the start of the cell cycle is dependent on Cln/CDK but independent of Whi5 (De Bruin *et al*, 2004) (unlike SBF expression, which is dependent on both). Furthermore, Wittenberg and Reed hypothesized that MBF activation may involve direct phosphorylation of Swi6 or phosphorylation of another MBF associated protein (Wittenberg & Reed, 2005). Our results show that MBF expression is indeed dependent on the action of the G1 cyclins, but is not abrogated by the loss of any single one. However, the expression timing is strongly influenced by the G1 cyclins, with the strongest effect caused by the loss of Cln3 (5-10 mins delay compared to the wild type, depending on the definition of targets, Supplementary S.5). In cells lacking Cln3, also the expression peaks of SBF and MBF are less delayed relative to each other. This more similar regulation of the two clusters in absence of Cln3 could be explained by a backup activation of MBF carried out by Cln1 and/or Cln2.

### Release from α-factor requires Cln2 and Cln3

The interference of α-factor synchronization with our knockout experiments first appeared to be a substantial drawback in our experimental setup. However, it allowed us to dissect an unexpected aspect of G1 cyclin action: their specific interaction with the pheromone-induced cell cycle arrest. We showed that cells lacking Cln2 as well as Cln3 are not able to exit from the arrest in a coordinated manner - resulting in a desynchronized population with no apparently oscillating genes. We can include *cln2*Δ*cln3*Δ to a previously described partially viable group of mutants (Barik *et al*, 2016), since we observed a five-fold higher fraction of dead cells after the release (∼25% compared to ∼5% in all other mutants (Supplement Figure S.9) indicating a severe effect of the deregulation. Mechanistically, the *cln2*Δ*cln3*Δ mutant failed to repress a class of genes linked to the pheromone response also after α-factor was no longer present in the medium.

G1 cyclins are known to compete with the pheromone response component Far1, a CDK inhibitor, by targeting it for degradation as their concentrations increase (Peter *et al*, 1993; McKinney *et al*, 1993). Far1 was initially shown to interact mainly with Cln2 (Chang & Herskowitz, 1990; Valdivieso *et al*, 1993), later Cln1 was proposed to have a comparable interaction (Oehlen & Cross, 1994). Since in our data the release defect only occurred in the *cln2*Δ*cln3*Δ mutant, we conclude that the interaction with the pheromone pathway can indeed not be completely independent of Cln1, since then also the *cln2*Δ single mutant would have difficulties in releasing. However, the effect of loosing Cln2 is much more severe, evident from the *cln1*Δ*cln3*Δ strain, which has no problems relieving the arrest although having a drastically longer G1 phase. In the *cln2*Δ*cln3*Δ mutant, Cln1 alone, without the strong initial “boost” by Cln3, does not sufficiently activate the positive feedback loop to reliably start G1 transcription and concurrently target Far1 for degradation. On the other hand, Cln2 alone (in the *cln1*Δ*cln3*Δ strain) can, following a lengthy G1 phase, eventually reach high enough concentrations to inactivate Far1 and initialize the cell cycle.

### Osmostress and α-factor arrest as a tools to increase effects of G1 mutations

In our experiments, the additional perturbations by osmotic stress and α-factor revealed further differences between the mutant strains that would not have been evident from unperturbed cultures. Both perturbations interfere with the cell cycle during G1 phase, which is also the phase most reliant on the action of the analyzed cyclins. Especially, the differences between Cln1 and Cln2 became evident in the stress experiments, for example in the quantitative analysis of the peak times (Figure 3B). While no effect of the loss of Cln1 alone was visible in the unstressed experiment, the global analysis of all oscillating genes in the stressed mutant showed a delay compared to the wild type, distinct from an even stronger effect in the Cln2 mutant.

Cell cycle synchronization with α-factor is frequently used in cell cycle studies, but is not without effect on the cells after the release (*e.g.* (Doncic *et al*, 2011; Eser *et al*, 2011) and this study). While here we could use this lasting perturbation to identify further differences between functions of the cyclins, some other effects might be masked by the initial decaying response to α-factor. For example, the only slightly altered length of the G1 phases of the Cln1 and Cln2 single and double mutants could be due to the stronger retainment of Whi5 in the nucleus in cells treated with α-factor (Doncic *et al*, 2011). The differences in Whi5 localization caused by the lack of Cln1 and Cln2, and therefore their potentially longer G1 phases would then not become visible. Accordingly, experimental setups utilizing α-factor synchronization should be handled with care, but can then still be very valuable for analyzing the cell cycle.

## Methods

### Strains

BY4741 (*MAT***a** his3Δ1 leu2Δ0 met15Δ0 ura3Δ0) haploid *S. cerevisiae* strain was used as parental strain. Deletion cassettes for all mutants were generated on a PCR-based protocol using pUG72 (Euroscarf accession number P30117) as template. Deletion cassettes were transformed into BY4741 and deletions were selected on minimal medium lacking uracil. Successful integration was controlled by PCR. The Ura3 selection marker was removed by expression of Cre recombinase from plasmid pSH68 (Euroscarf accession number P30674).

### Growth conditions and synchronization

All experiments were performed in SD medium. For osmostress experiments, 0.4 M NaCl was added to SD medium. For time course experiments, cells were synchronized as described before (Futcher, 1999) with small modifications. Cells were grown over night and inoculated in fresh medium to OD_600_ = 0,05 and grown till OD_600_=0,2. Afterwards, cells were washed, re-suspended in the same volume of fresh media and 5 μg/ml alpha factor. After 3 hours of synchronization, cell were washed and released in fresh medium, containing 0.4 M NaCl for osmostress experiments.

### Time course, RNA extraction and sample preparation

Samples were taken over 150 min (60 min every 5 min, afterwards every 10 min) and frozen in liquid nitrogen. RNA was extracted with Nucleospin 96 RNA Kit (Machery-Nagel, cat 740466.4) with small modifications. Cell lysis was done for 30 min at 30^∘^C by adding 450μl of lysis buffer containing 1 M sorbitol, 100 mM EDTA, and 0.45 ml lyticase (10 IU/ml). The rest of the RNA extraction was performed according to manufacturer’s details. Extracted RNA was converted to cDNA, barcoded and sequenced with Illumina HiSeq 2500.

### RNA sequencing, data processing pipeline

RNA reads were aligned to reference genome using BOWTIE and filtered for rRNAs. Every sequence was normalized for PCR bias using UMIs (Kivioja *et al*, 2011) and cleaned if the reads align more than once to the genome. Reads were normalized to total expression reads, which was defined as 10^6^ and genes with expression below 10 reads were excluded from the analysis. Time points were removed if total reads were less than 7⋅10^4^ (Supplement Figure S10). All experiments were done in replicates. Since the correlation between replicates was very high (Supplement Figure S11) we merged the experiments and worked with the mean of expression.

### Clustering and functional enrichment of clusters

The changes in mRNA expression in the wild type over time were clustered once for unstressed conditions and once for osmostress by k-means clustering to group similar dynamics together. The same genes were plotted for the mutants and expression was compared to wild type. Clusters were tested for functional enrichment using GO enrichment provided by YeastMine database (Cherry *et al*, 2012) and via mapping to a generic proteomap (Liebermeister *et al*, 2014), which sorts proteins (here: transcripts) into functional groups.

### Detection of periodic genes

The MoPS package (Eser *et al*, 2014) for R was used and extended to quantify periodicity for individual genes. In addition to the periodicity score calculated by the package, we calculated the Bayesian Information Criterion (BIC) to punish good fits with low numbers of measurement points since the MoPS periodicity score does not explicitly account for the number of data points for each gene. We neglected genes with less than 13 valid time points. The shaping parameter of the MoPS package was not used since it causes many false positives. As in Eser *et al*. (Eser *et al*, 2014), we defined a cutoff value for the periodicity score based on the result of the 200 most periodic genes listed in cyclebase (Santos *et al*, 2014) (20% false negative rate). We used the same approach for the BIC normalized by the estimated amplitude of each gene, to select only genes with a sufficiently good model fit as periodic (Supplementary S.2).

### Local regression of peak time shifts

Average behavior of peak time shifts in the mutants was visualized using local regression (locally weighted scatterplot smoothing, lowess) of the estimated wild type peak times scattered against the peak times of the stressed and unstressed mutants. The matlab function smooth with the option ‘rlowess’ was used, performing robust local regression with a first degree polynomial using a span of 20% of the total data points.

### Transcription factor network analyses

We obtained a generic yeast transcription factor (TF) network from the YEASTRACT database (Teixeira *et al*, 2014), including all TF-target pairs that were shown to interact by DNA binding and expression evidence (8685 edges, Supplementary file 3). Target gene groups’ behavior was again analyzed with MoPS to score periodicity of the gene expression driven by a specific transcription factor.

### Automatic detection of transcription factor target peak expression

Expression trajectories of all target genes of a selected transcription factor were averaged and smoothed with a spline interpolant (matlab interp1 function with option ‘spline’, final resolution of 1 minute). A simple peak detection was implemented by searching for points that are expressed higher than the next and previous 10 points (’detected peaks’). Thus detected peaks were only considered if they occured after all peaks of the target genes assigned to the previous cell cycle phase (’valid peaks’). For cases were several transcription factors define one phase transition, valid peaks’ times were averaged to obtain the phase transition time.

### Quantification of delays via cross correlation

Cross correlations were calculated with the MatLab function ‘crosscorr’ between the pairs of mean target expression (MBF and SBF in one mutant or MBF in a mutant and MBF in the wild type). To obtain mean target expression, each gene trajectory was first normalized to its temporal mean, linearly interpolated to 1 min time steps, then the trajectories of all target genes of MBF or SBF (defined from different sources, Supplementary File 4) were averaged.

### Microfluidics, microscopy and image analysis

To control synchronization and release from α-factor arrest, CellASIC ONIX Microfluidic Platform in combination with Y04C for haploid cell was used according to manufacturer’s instructions. A mixed culture with OD_600_=0.4 was loaded to the plates incubated at 30^∘^C and exposed to 5 μg/ml α-factor in SD for 3 h with a pressure of 2 psi. Afterwards α-factor was removed by washing plates with SD for 10 min with a pressure of 5 psi. Imaging was performed every 10 min for 7h using a Olympus IX83 inverted microscope with a motorized stage. Brightfield images were acquired with an iXon EMCCD Cameras, ANDOR. Detection of bud emergence was performed manually utilizing BudJ: an ImageJ plugin to analyze fluorescence microscopy images of budding yeast cells (http://www.ibmb.csic.es/groups/spatial-control-of-cell-cycle-entry).

**Table 1:** Transcription factors with oscillating target expression. The lower factors are not shown in Figure 4 if they have less than 3 target genes (f) or are not connected to any other transcription (u).

| ID | ORF | MoPS score | MoPS $\phi$ | # targets | short description (SGD) | why not in network |
| --- | --- | --- | --- | --- | --- | --- |
| DIG1 | YPL049C | 0.81 | 0 | 6 | MAP kinase-responsive inhibitor of the Ste12p transcription factor |  |
| PHD1 | YKL043W | 1.77 | 3 | 32 | Transcriptional activator that enhances pseudohyphal growth |  |
| SOK2 | YMR016C | 1.06 | 3 | 375 | Nuclear protein that negatively regulates pseudohyphal differentiation |  |
| MSN2 | YMR037C | 0.93 | 5 | 693 | Stress-responsive transcriptional activator |  |
| PHO4 | YFR034C | 1.27 | 5 | 65 | Basic helix-loop-helix (bHLH) transcription factor of the myc-family |  |
| TEC1 | YBR083W | 1.34 | 5 | 309 | Transcription factor targeting filamentation genes and Ty1 expression |  |
| STE12 | YHR084W | 1.26 | 6 | 937 | Transcription factor that is activated by a MAPK signaling cascade |  |
| MSS11 | YMR164C | 1.07 | 7 | 9 | Transcription factor |  |
| TUP1 | YCR084C | 0.81 | 7 | 109 | General repressor of transcription |  |
| CIN5 | YOR028C | 0.92 | 8 | 201 | Basic leucine zipper (bZIP) transcription factor of the yAP-1 family |  |
| MBP1 | YDL056W | 1.66 | 8 | 60 | Transcription factor |  |
| ASH1 | YKL185W | 1.04 | 9 | 64 | Component of the Rpd3L histone deacetylase complex |  |
| MGA2 | YIR033W | 1.04 | 12 | 13 | ER membrane protein involved in regulation of OLE1 transcription |  |
| SWI4 | YER111C | 2.14 | 12 | 134 | DNA binding component of the SBF complex (Swi4p-Swi6p) |  |
| HAC1 | YFL031W | 1.2 | 13 | 9 | Basic leucine zipper (bZIP) transcription factor (ATF/CREB1 homolog) |  |
| SWI6 | YLR182W | 2.09 | 13 | 41 | Transcription cofactor |  |
| HIR2 | YOR038C | 1.71 | 22 | 8 | Subunit of HIR nucleosome assembly complex |  |
| HCM1 | YCR065W | 1.16 | 24 | 10 | Forkhead transcription factor |  |
| FKH2 | YNL068C | 1.12 | 33 | 23 | Forkhead family transcription factor |  |
| HMS1 | YOR032C | 0.79 | 44 | 7 | bHLH protein with similarity to myc-family transcription factors |  |
| MIG2 | YGL209W | 1.04 | 46 | 4 | Zinc finger transcriptional repressor |  |
| MCM1 | YMR043W | 2.17 | 55 | 115 | Transcription factor |  |
| YOX1 | YML027W | 1.31 | 59 | 97 | Homeobox transcriptional repressor |  |
| SWI5 | YDR146C | 1.13 | 61 | 83 | Transcription factor that recruits Mediator and Swi/Snf complexes |  |
| ACE2 | YLR131C | 1.08 | 62 | 116 | Transcription factor required for septum destruction after cytokinesis |  |
| YHP1 | YDR451C | 1.21 | 62 | 38 | Homeobox transcriptional repressor |  |
| FLO8 | YER109C | 0.94 | 63 | 37 | Transcription factor |  |
| CUP9 | YPL177C | 0.78 | 65 | 16 | Homeodomain-containing transcriptional repressor |  |
| HIR1 | YBL008W | 1.5 | 20 | 1 | Subunit of the HIR complex | f |
| YRR1 | YOR162C | 0.88 | 22 | 2 | Zn2-Cys6 zinc-finger transcription factor | f |
| AZF1 | YOR113W | 1 | 52 | 2 | Zinc-finger transcription factor | f |
| RLM1 | YPL089C | 1.28 | 3 | 34 | MADS-box transcription factor | u |
| MAL33 | YBR297W | 1.33 | 10 | 11 | MAL-activator protein | u |
| UGA3 | YDL170W | 2.11 | 13 | 4 | Transcriptional activator for GABA-dependent induction of GABA genes | u |
| PIP2 | YOR363C | 1.01 | 63 | 44 | Autoregulatory, oleate-activated transcription factor | u |
| HAL9 | YOL089C | 0.88 | 69 | 8 | Putative transcription factor containing a zinc finger | u |

**Table 2:** Overview of selected transcription factors for definition of cell cycle phase duration.

| Group | Transcription factor | Periodicity score WT | Estimated target peak time WT [min] | Event at peak (from literature) | References |
| --- | --- | --- | --- | --- | --- |
| I | Swi4 | 2.14 | 12 | G1/S transition | McIntosh *et al*, 2000; Iyer *et al*, 2001; Simon *et al*, 2001; Lee *et al*, 2002; Teixeira *et al*, 2006; Ferrezuelo *et al*, 2009 |
|  | Swi6 | 2.09 | 13 |  |  |
|  | Mbp1 | 1.66 | 8 |  |  |
| II | Hir1 | 1.5 | 20 | S/G2 transition | Sherwood *et al*, 1993; Spector *et al*, 1997 |
|  | Hir2 | 1.71 | 22 |  |  |
| III | Mcm1 | 2.17 | 55 | G2/M transition | Loy *et al*, 1999; Kumar *et al*, 2000 |
| IV | Swi5 | 1.13 | 61 | Mitotic exit, reset for new G1 | Dohrmann *et al*, 1992, 1996; McBride *et al*, 1999; Laabs *et al*, 2003 |
|  | Ace2 | 1.08 | 61 |  |  |

**Table 3:**
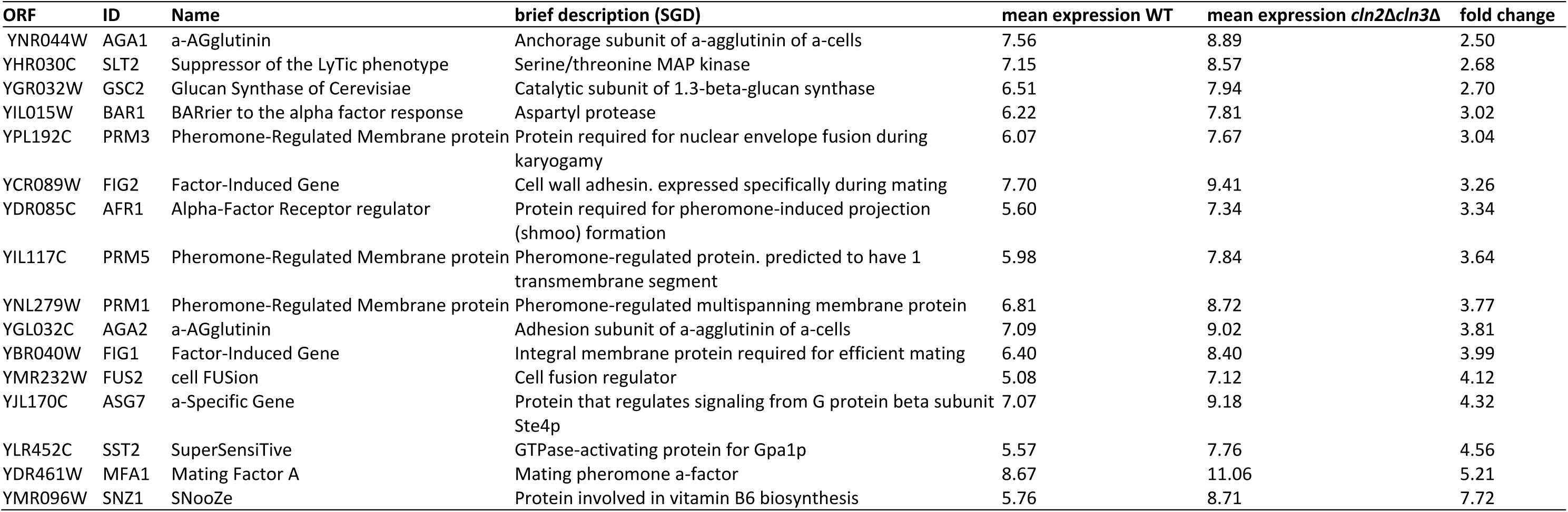
Group of α-factor induced genes with higher expression over the experiment in *cln2Δcln3Δ* (also see Figure 6)

## Acknowledgments

This work was supported by the PhD program of the Max-Delbrück-Center for Molecular Medicine (LT), Deutscher Akademische Austauschdienst (DAAD, LT) and by the German Excellence Initiative (Caroline-von-Humboldt professorship to EK).

## Supplementary Figures

**Figure S1: Overview of differentially expressed gene clusters** found by k-mean clustering on wild type (WT) gene expression trajectories under unstressed growth conditions. Compared to all strains (upper panel represent single deletions and lower panel represents double mutants). Mean expression is shown for Wild type in gray and corresponding genes in the mutants strains are plotted in colors

**Figure S2: Calculate MoPS periodicity score and BIC.** Blue dots are all yeast genes, orange ones the top 200 oscillating genes from cyclebase. Dotted lines denote the 20\% quantiles of the period genes’ distributions and was used as cut-off value.

**Figure S3: Comparison of the found sets of oscillating genes. (A)** Genes in the 5 oscillating clusters (Figure 2) and oscillating genes identified by MoPS (Eser *et al*, 2014). **(B)** MoPS list compared to previously defined oscillating gene sets (the 500 most oscillating genes from cyclebase (Santos *et al*, 2014) and from Spellman *et al*. (Spellman *et al*, 1998). Visualization by BioVenn (Hulsen *et al*, 2008).

**Figure S4: Target expression patterns** of selected transcription factors and cell cycle phase classification in all experimental conditions (as shown in detail for the wild type in Figure 4)

**Figure S5: Delays in expression based on different target gene lists for SBF and MBF.** Analysis is the same as in the main Text Figure 5, with target gene lists as defined by Eser *et al*. (Eser *et al*, 2011), MacIsaac *et al*. (Macisaac *et al*, 2006) and Ferrezuelo *et al*. (Ferrezuelo *et al*, 2009).

**Figure S6: Acute osmotic stress response. (A)** Wild type expression of stress responsive gene clusters (upper panel, line represents mean, shaded area 25% and 75% quantiles), annotated with cell cycle phases. The lower panel shows the mean expression of the corresponding gene clusters in the mutant strains. The wild type behavior is plotted for comparison. **(B)** Functional classification based on a proteomap (Liebermeister *et al*, 2014), genes are colored according to the stress cluster in A.

**Figure S7: Timing of budding after α-factor synchronization release** in microfluidic setup for wild type and all deletion strains. Cells were synchronized for 180 min (red column), at time point 0 **α**-factor washed-out. Several colonies were imaged every 10 min and newly occurred buds were counted for each image (blue histogram), red line represent colony size as total number of cells and buds. Budding was detected for two cell cycles after the release.

**Figure S8: Number of differentially expressed genes in the mutants**, normalized to the defined cell cycle phases. Expression trajectories of the mutants were normalized by scaling corresponding phases to the wild type (WT) phase durations. Samples taken at switching times between the phases were assigned to the previous phase. To calculate differential expression, WT expression was linearly interpolated at the now normalized mutant time points. Genes were then deemed differentially expressed if at least *n* ρ = 3 time points were changed more than 3.36-fold (log_2_(1.75)) or a low p-value in a Kolmogorov-Smirnow-test compared to the WT expression (ρ<10^-8^ for *n* _ρ_ = 0, ρ<8×10^-5^ for *n* _ρ_ =1 and ρ<0.002 for *n* _ρ_ = 2).

**Figure S9: Viability test,** cells were stained with propidium iodide and percentage of viable cells were measured using Flow cytometry. Cells were synchronized (as described in methods) and sample were taken every 15 min over 150 min. Since no differences in viability over time were detected, mean viability in percent and standard deviation for each strain are depict.

**Figure S10: Statistics of total reads per sample.** Samples with less than 7×10^4^ reads were excluded from the analysis (blue fields in the heatmap plot, red line in histogram). Note that in the histogram missing samples are counted in the first bin, whereas they are specifically shown in gray in the heatmap plot.

**Figure S11: Mean correlation of the RNAseq replicates.** The Pearson correlation coefficient was calculated for the expression value of all timepoints of a specific gene between two replicates and subsequently averaged over all timepoints for this specific pair of samples. The boxplots show the distribution of all such average correlation coefficients of which the replicate stated on the x axis occurs, and hence the overall correlation of each replicate with all others.

